# Intranasal inoculation of *Cryptococcus neoformans* in mice produces nasal infection with rapid brain dissemination

**DOI:** 10.1101/709204

**Authors:** Carolina Coelho, Emma Camacho, Antonio Salas, Alexandre Alanio, Arturo Casadevall

## Abstract

*Cryptococcus neoformans* is an important fungal pathogen, causing life-threatening pneumonia and meningoencephalitis. Brain dissemination of *C. neoformans* is thought to be a consequence of an active infection in the lung which then extravasates to other sites. Brain invasion results from dissemination via the bloodstream, either by free yeast cells in bloodstream or Trojan horse transport within mononuclear phagocytes. We assessed brain dissemination in three mouse models of infection: intravenous, intratracheal, and intranasal. All three modes of infection resulted in dissemination of *C. neoformans* to the brain in under 3 hours. Further, *C. neoformans* was detected in the entirety of the upper respiratory tract and the ear canals of mice. In recent years, intranasal infection has become a popular mechanism to induce pulmonary infection because it avoids surgery but our findings show that instillation of *C. neoformans* produces cryptococcal nasal infection. These findings imply that immunological studies using intranasal infection should assume the initial sites of infection of infection are brain, lung and upper respiratory tract, including the nasal airways.

**Importance:** *Cryptococcus neoformans* causes an estimated 181, 000 deaths each year, mostly associated with untreated HIV/AIDS. *C. neoformans* has a ubiquitous worldwide distribution. Humans become infected from exposure to environmental sources and the fungus lays dormant within the human body. Upon immunosuppression, such as AIDS or therapy-induced as required by organ transplant recipients or autoimmune disease patients, cryptococcal disease reactivates and causes life-threatening meningitis and pneumonia. This study has detected that upon contact with the host, *C. neoformans* can quickly (a few hours) reach the host brain and will also colonize the nose of infected animals. Therefore, this work paves the way to better knowledge of how *C. neoformans* travels through the host body. Understanding how *C. neoformans* infects, disseminates and survives within the host is critically required so that we can prevent infections and the disease caused by this deadly fungus.

## Introduction

The genus *Cryptococcus* is populated by environmental fungi, wood-rotting fungi most commonly associated with trees but also with bird guano. Two species of *Cryptococcus* can cause life-threatening disease in humans, characterized mainly by pneumonia and life-threatening meningoenchephalitis. *C. neoformans* var. *grubii* is the most prevalent pathogen of the genus *Cryptococcus*, causing approximately 200,000 deaths each year, mostly associated with HIV-positive individuals, while the closely related species *C. gattii* was responsible for an outbreak in British Columbia in apparently immunocompetent individuals (1–3).

The current paradigm for the pathogenesis of human cryptococcosis emerged from a series of observations made over several decades. Exposure is due to inhalation of infectious propagules from the environmental niches. Colonization of the lungs by spores, desiccated yeasts or yeast cells is thought to be quickly controlled by the human immune system via granuloma formation (called cryptococcoma). In the 1950s, silent cryptococcal granulomas were reported in lungs, establishing a parallel to latent tuberculosis (4). In support of this notion, serological studies have established that adults have antibodies to *C. neoformans* (5) or had delayed hypersensitivity skin reactions (6), consistent with asymptomatic infection. Both teenagers or children under the age of 5 living in urban areas are immunoreactive against *Cryptococcus* (7, 8). Finally HIV-patients are infected with a serotype most commonly isolated from their place of birth or where they spent their infancy (9). Hence, humans are exposed very early in life to this yeast without developing noticeable disease. Silent or latent infections due to containment in the lungs is further supported by the observation that some recipients of lung transplants develop a cryptococcal infection originated from the donor lung (10–12). Even though most infections of *C. neoformans* in humans are asymptomatic, a significant and unknown proportion of humans become latent carriers of *C. neoformans* which may reactivate as host immunity declines.

Given that initial human infection is thought to start from the lungs, research in pathogenesis of *C. neoformans* commonly uses pulmonary infection models. The two inoculation routes most commonly used are intratracheal and intranasal infection. Intratracheal infection delivers *C. neoformans* directly to the respiratory tree but requires surgery and significant skill. In contrast, intranasal infection is done by depositing a solution containing *C. neoformans* in the nose of an anesthetized mice, which then inhales it as these animals are obligate nose breathers. Since intranasal infection does not require surgery, it has become a popular mode of inducing *C. neoformans* infection in mice. Intranasal infection has been accepted as a procedure for inducing pulmonary infection without much investigation as to what actually happens after nose deposition of an infective inoculum. In some cases researchers use a third route of infection, the intravenous route, since it allows a rapid dissemination to the brain, presumably by bypassing lung immune response (13, 14). Intravenous model in outbred mice have been shown to be a relevant model when compared to human infection (15–17).

In this study we compared intravenous, intratracheal and intranasal infection and observed that all produce rapid brain dissemination. Additionally, we detected that intranasal inoculation leads to presence of yeasts in upper respiratory airways and auditory tract. These observations have important implications for the interpretation of intranasal studies when considering immune responses and aspects of pathogenesis.

## Materials and Methods

*C. neoformans* was grown from frozen glycerol stocks into a Yeast Peptone Dextrose (YPD) plate for 2 days, and then cultured overnight at 30 ^o^C in YPD broth with shaking. We used a strain from H99E lineage (available from Lodge laboratory), the H99O strain, a close relative of the original isolate of H99 (18), and the R265 strain of *C. gattii*, obtained from American Type Culture Collection.

C57BL/6J mice, aged 8–10 weeks, were obtained from Jackson Laboratories and infected with the indicated 5×10^3^ (low dose) to 5×10^5^ CFU (high dose) in a final volume of 40 μl of sterile PBS (19). Intravenous (IV) injections were performed by 40 μl injection in the retroorbital sinus of the animal under isoflurane anesthesia. Intranasal (IN) experiments were performed by placing 40 μl of yeast suspension into the mouse nares while under isoflurane anesthesia (20). Intratracheal (IT) infections were performed under xylazine-ketamine anesthesia. Animal’s neck was exposed, the trachea was exposed via midline incision and yeasts inoculated with a 25G gauge syringe directly into the trachea. Incision was closed with Vetbond (3M, St. Paul, MN, USA). Mice were monitored daily for signs of stress and deterioration of health throughout the experiment. All animal experiments were approved by Johns Hopkins University IACUC under protocol number MO18H152.

To measure fungal burden, mice were euthanized, exsanguinated via terminal retroorbital bleeding, tissues removed and macerated by passing through a 100 μm mesh into sterile PBS. The tissue homogenate was then plated into YPD agar plates, and colony forming units (CFU) were quantified. For one experiment we rinsed the brains (to remove possible yeast contaminations from the exterior surface of the brain during necropsy). In one experiment we included a non-infected mouse (“sentinel”) to test for accidental contamination of tools and materials during necropsy.

For histological analysis of the skull, the skin, lower jaw and tongue were removed and remainder of skull was fixed in Formacal and processed for routine histology. A sagittal cut was performed through the middle section of the mouse skull and consecutives 4 μm sections, with 40 μm intervals were cut for Hematoxylin-Eosin, Mucicarmine, Grocott-Methenamine Silver and immunofluorescence. For immunofluorescence 18B7 monoclonal antibody against capsular polysaccharides (21) was used and then detected with anti-mouse IgG_1_-Alexa 488 conjugate antibody. Hematoxylin & eosin sections were scanned using a slide scanner at the Oncology Tissue Services Core (School of Medicine, Johns Hopkins University). Remaining sections were imaged using an Olympus AX70 microscope (Olympus America, NY, USA). Image cropping and annotation were performed using ImageJ (22).

## Results

We compared 3 routes of infection (IV, IN, and IT) to ascertain the kinetics of *C. neoformans* dissemination to the brain (Fig.1). We had previously verified the infectious dose of 5×10 ^5^ CFU induces 100% mortality in 22 days for H99E. We quantified fungal burden in blood, lung and brains. In two mice, we detected fungi in blood although in these we also detected bacteria in blood (data not shown). This suggests that animals with *C. neoformans* in the blood had severe systemic infection that may have predisposed to concomitant bacterial infection. We conclude that the majority of animals clear *C. neoformans* from the bloodstream. After 7 days of infection, we measured increased numbers of yeast cells in the brain after IV inoculation (more than 10^5^, upper limit of detection) than after IN or IT inoculation (below 10^4^ CFU). In the lung there was a slight increase in fungal burden when the IT route is used, as compared to the IV route. We did not detect differences in brain fungal burden between IN and IT inoculation. Remarkably, we noted there was a substantial amount of yeast in mouse brains as early as day 3 after inoculation of yeast by both IN or IT, which prompted us to ask how quickly *C. neoformans* disseminates to the mouse brain.

**Figure 1.**
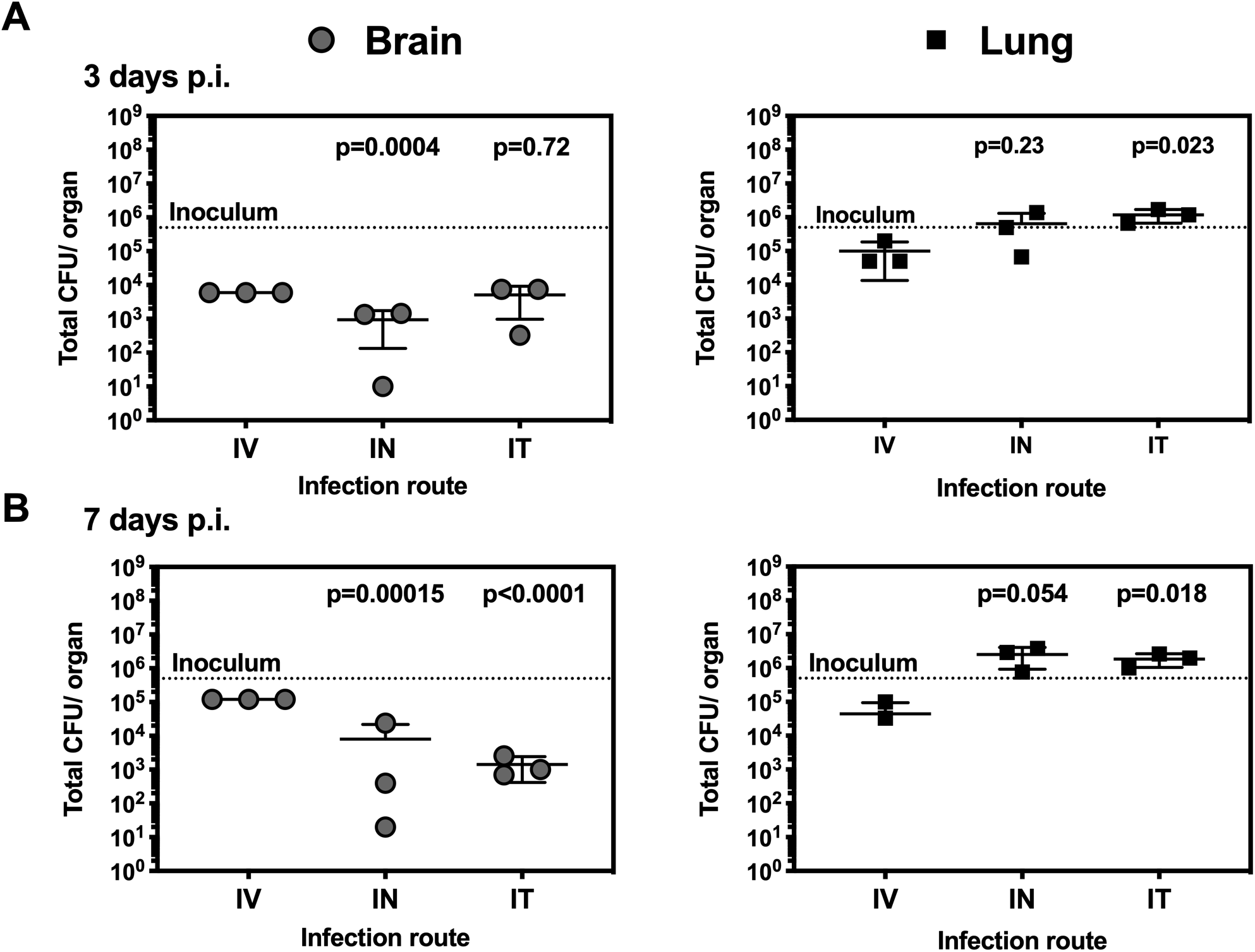
*C. neoformans* dissemination to the brain and lung after different inoculation routes. Mice were infected with 5 x 10^5^ H99E strain of *C. neoformans* via intravenous (IV), intranasal (IN) and intratracheal (IT) route. Colony forming units (CFU) analyzed at A) 3 days post infection (dpi) and, B) 7 dpi. Each data point represents one individual mouse and bars represent mean and SD. Numbers represent P-values calculated from two-stage linear step-up procedure of Benjamini, Krieger and Yekutieli, with Q = 5%, as compared to IV infection.

We focused on the intranasal model since this procedure is non-invasive, requiring only brief anesthesia and no surgery. We found that 3 h post infection with 5×10^5^ yeast inoculum, the fungal burden in the brain was in the order of hundreds of CFU for most animals (Fig. 2A). The surprising finding that yeasts were detectable in the brain within hours after IN inoculation prompted us to perform additional controls. For one experiment we rinsed the brains (to remove possible yeast contaminations from the exterior surface of the brain during necropsy) but we got comparable results to the first experiment (data pooled from both experiments is shown). We also culled one non-infected mouse (“sentinel”) at the end of the experiment to measure eventual contamination of tools and materials during necropsy that could cross-contaminate the media plates. We recovered no CFU from the sentinel mouse which gave us confidence that the fungal burden we detect are result from brain infection and not accidental contamination (data not shown). To investigate if early dissemination could occur at lower doses we infected animals with 5×10^3^ CFU (at this dose *C. neoformans* causes 80% mortality at 40 days). We still observed quick dissemination to the brain, indicating that this was not a consequence of exposure to overwhelmingly high numbers of yeasts. Finally, we observed quick dissemination to the brain with strain H99O, a strain closely related to the original H99 clinical isolate of *C. neoformans*, as well as the R265 strain of *C. gattti*, the strain associated with the Vancouver outbreak (Fig. 2B). It is possible that the detected yeast burden is arrested in the small capillaries (23–26), or in the post-capillary venules (27) and has not yet invaded the brain parenchyma. However, this is consistent with invasion of the brain since upon arrest in capillaries, *C. neoformans* quickly crosses endothelial barriers (24) and possibly the blood brain-barrier. These experiments show that after intranasal infection, pathogenic strains of *C. neoformans* and *C. gattii* disseminate to the mouse brain in as little as 3 h.

**Figure 2.**
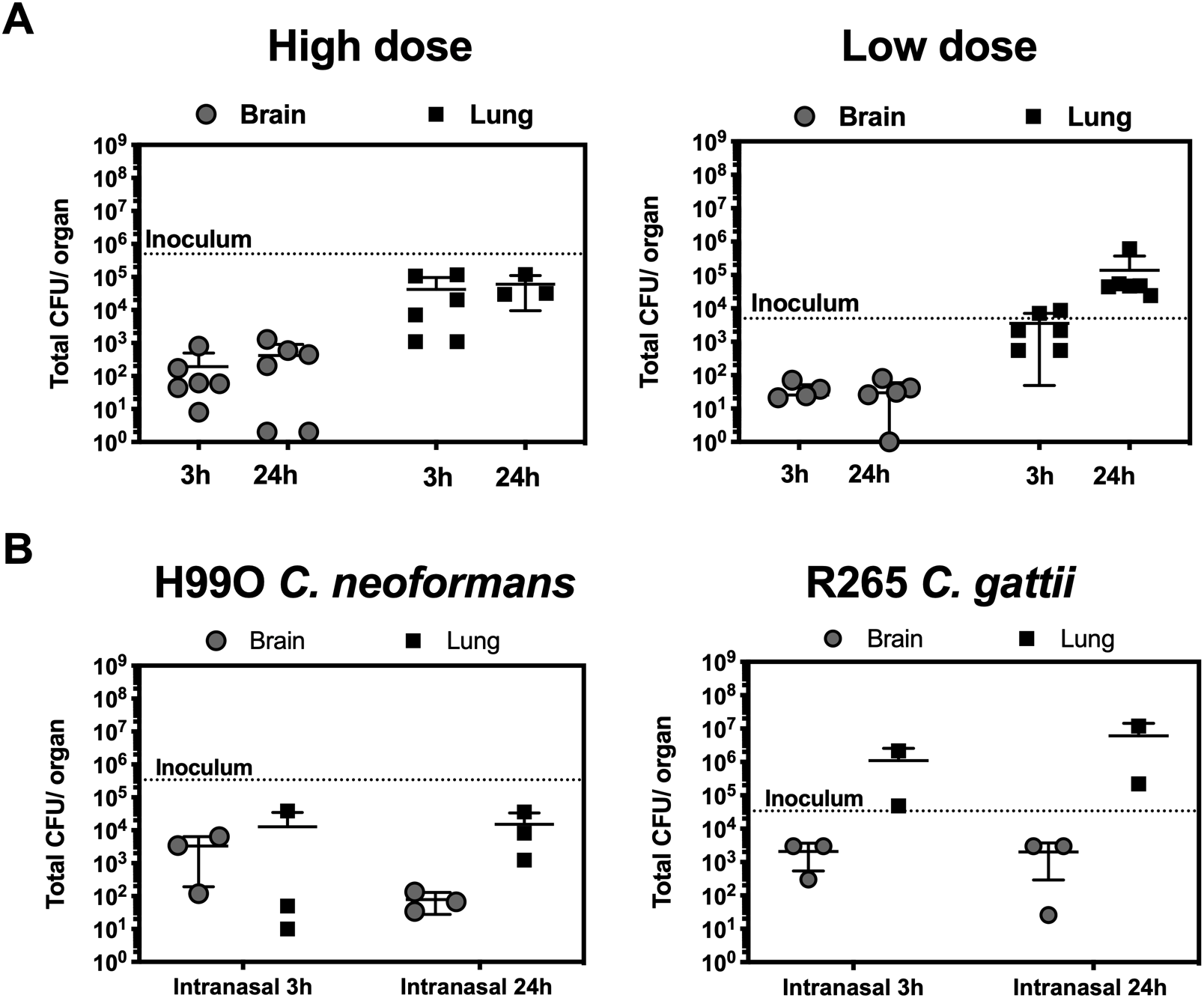
*C. neoformans* is detected in the brain as early as 3 h post infection (hpi) after IN infection. A) Mice were infected with either 5 x 10^5^ CFU (High dose) or 5 x 10^3^ CFU (Low dose) of H99E via intranasal instillation and euthanized at the indicated time to measure yeast tissue burden. B) Mice were infected with H99O strain of *C. neoformans* or R265 strain of *C. gattii*. Each data point represents one individual mouse and bars represent mean and SD.

To confirm presence of yeast in the brain, we performed immunofluorescence in skulls of infected mice, using a monoclonal antibody against *C. neoformans* capsular GXM (21). In agreement with the results of CFU quantification, we detected yeast cells (positive immunostaining and with typical morphology) in the mouse brain at 24 h post infection (Fig. 3A, inset). We wondered if it was possible for *C. neoformans* could provoke damage to the nose mucosa and from there access the brain through damaging the olfactory mucosa and using the olfactory nerves to traverse the cribriform plate (as shown for bacterial pathogens (28, 29) or if after damage the mucosa access the host bloodstream for dissemination. We reasoned that if dissemination occurred through the olfactory system, then the olfactory bulb would contain the majority of yeasts, compared to the remainder of the brain. We infected mice and separated their olfactory bulb from the remainder of the brain (Fig. 3B). We found that the olfactory bulb contains similar amounts of yeasts to the remainder of the brain, despite its small size compared to the rest of the brain.

**Figure 3.**
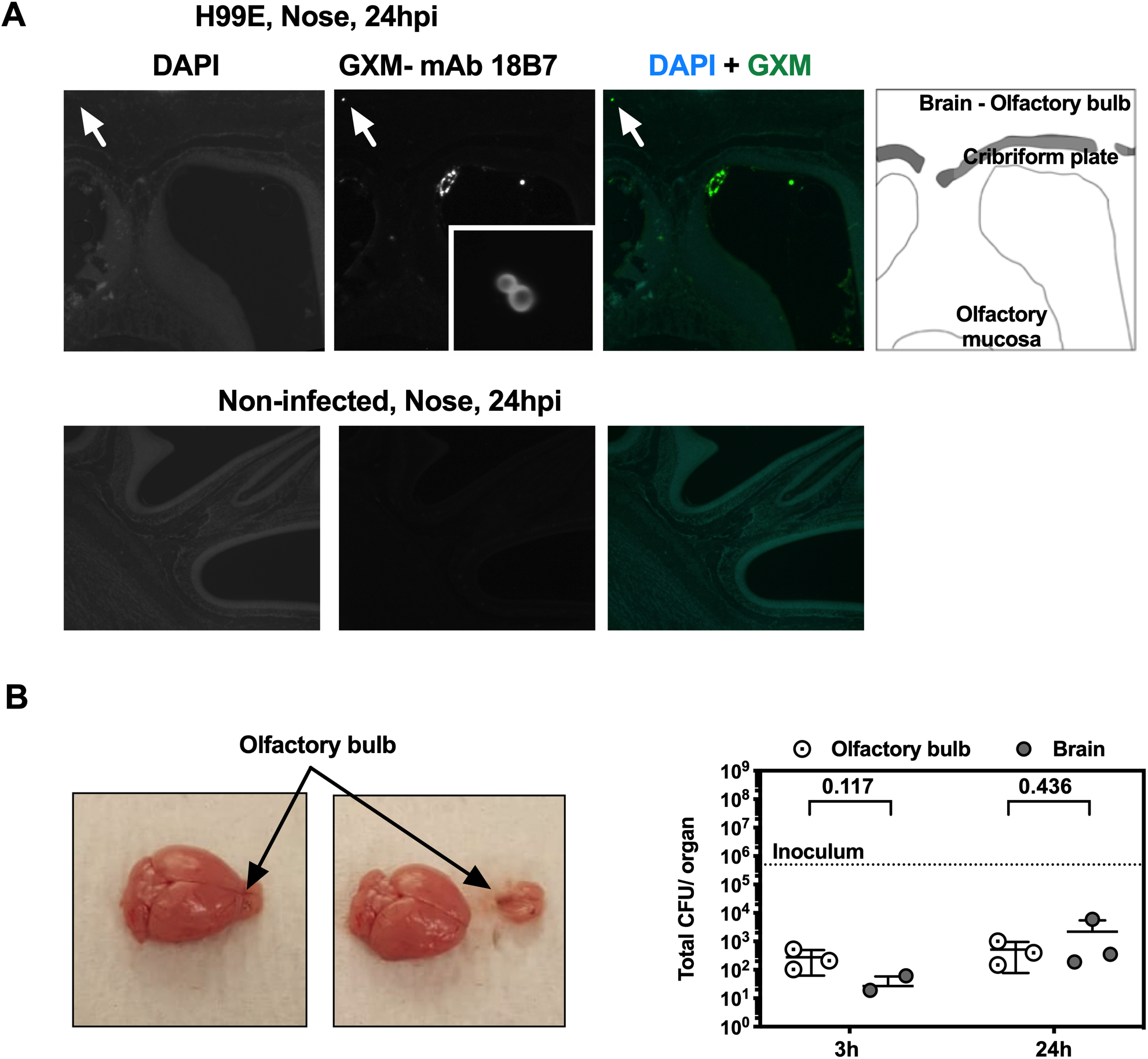
*C. neoformans* is detected in the brain and the olfactory bulb as early as 3 h post infection (hpi) after IN infection. A) Mice were infected with 5 x 10^5^ CFU of *C. neoformans* via intranasal instillation, euthanized at the indicated time, and processed for histology. Yeast cells could be observed in the olfactory bulb of the brain of mice at 24 hpi (top panel, arrows). Non-infected animals were processed under the same conditions to show specificity of staining of yeast capsular immunofluorescence (bottom panel). B) Mice were infected with high dose of *C. neoformans*. At the indicated hpi, olfactory bulb was separated from the remainder of the brain (illustrated in right panel) and CFU were counted (left panel). Each data point represents one individual mouse and bars represent mean and SD. Numbers represent P-values calculated from two-stage linear step-up procedure of Benjamini, Krieger and Yekutieli, with Q = 5%.

To investigate the interaction of *C. neoformans* with the nose, and possible damage to the nose mucosa that could facilitate dissemination, we performed histological studies using a combination of histology, immunofluorescence, Grocott-Merwald-Silver, and mucicarmine (Fig. 4). We found yeasts scattered throughout the entire upper respiratory tract, particularly in the turbinates of the nose, and some yeasts close to the cribriform plate (Fig.4A-C), as well as in the ear canals of mice (Fig. 4D). Yeasts were abundant in airways, surrounded with a material that is likely mucus secretions from the host, but that stained abundantly with capsular GXM antibody, indicating a component of secreted polysaccharide in the airways (Fig. 4B). Budding forms rested on the respiratory epithelium cilia and olfactory epithelium layer, and yeast numbers increased in number from 3 h to the 24 h post infection and increased immunostaining of GXM (Fig. 4F). We noted at 24 h post infection there were rare enlarged yeast cells within the nose turbinates and ears (Fig. 4C-G) whose cell body diameter is above 10 μm, and therefore can be consider titan cells (30–32). The finding of titan cells in the nose of mice is concordant with Lima *et al*., (33) which found enlarged cells in noses of guinea pigs after some days of infection. Histopathology analysis detected no damage to the host epithelial layers as well as no inflammatory infiltrate from 3 h to 24 h post infection; it seems that either yeasts go undetected or they do not cause enough tissue damage to trigger inflammation in murine host in the early stages of infection. We conclude that, up to 24 h post-infection, presence of *C. neoformans* in nose of mice causes minimal, if any, damage to nose mucosa and the nose shows no sign of inflammation at this stage of infection.

**Figure 4.**
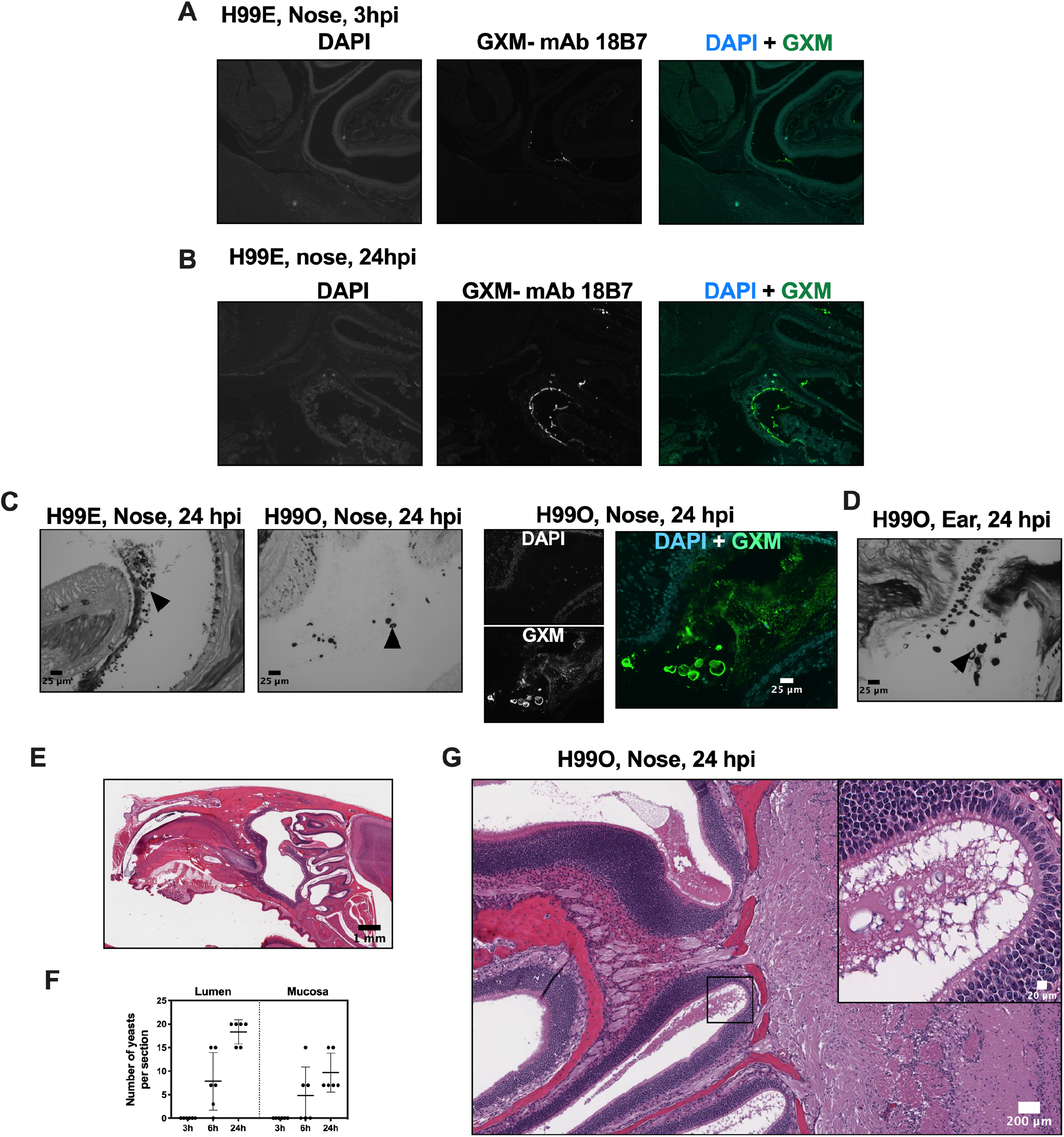
Histopathology of mouse nasal cavities upon *C. neoformans* infection. Colonization and secretion of GXM in airways of mice. Enlarged yeasts could be observed in airways and in ears. Mice were infected with 5 x 10^5^ CFU of H99E or H99O strains (as indicated) of *C. neoformans* via intranasal instillation, euthanized at the indicated time and processed for histology. Immunofluorescence staining of medial sagittal cuts of mouse skulls after staining with DAPI nuclear counterstain and anti-GXM-monoclonal antibody 18B7. A) Aspects of the cribriform plate and the nose turbinates showing yeasts in the airway. B) Accumulation of GXM in the nose mucosa. Specificity of staining was confirmed by performing immunostaining in a non-infected mouse (not shown); C) Grocott-Methenamine Silver staining (left, middle) and immunofluorescence (right panels) showing enlarged yeast cells in nose airways (arrowheads). D) Grocott-Methenamine Silver staining showing enlarged yeast cells in auditory canal (arrowhead, left panel) as well as abundant yeasts in auditory canals (right panel). Scale bars 25 μm. E) Hematoxylin-eosin staining of entire skull (left) (3 h post-infection). F) quantification of yeasts in nose and yeasts in airways. G) Aspect of nose mucosa, close to cribriform plate (24h post-infection), displaying lack of inflammation and showing enlarged yeast cells (inset).

## Discussion

Animal models of infection are critical tools for understanding the pathogenesis of infectious diseases. In the cryptococcal field, mice are the most commonly used mammalian species to study pathogenesis and immune responses. We compared the kinetics of brain dissemination in three models of cryptococcal infection that are commonly used: IV, IN and IT. We were particularly interested in IN infection since this approach has become increasingly popular in the field. We found early dissemination to the brain and colonization of upper respiratory tract by *C. neoformans* and *C. gattii*. Our data suggest early invasion of the brain and a colonization of the nose by both cryptococcal species in mouse models.

Quick dissemination to the brain observed in our mouse model is compatible with an hematogenous route of dissemination. Recent work has proposed that escape from the lungs occurs due to phagocyte drainage through the lymphatic system (34). It has not yet been shown how *C. neoformans* escapes from the lymph nodes into the bloodstream (35–37), but this will likely be elucidated in the future. We detected no yeasts in the circulating blood of mice, which raises a question of how the bloodstream is so quickly invaded and quickly cleared. Other groups observed that in IV infection fungi are cleared from the bloodstream in hours or the day after infection (27, 35), with fungemia only resurfacing very late in the infection. These observations, together with ours, imply that yeast transit time in blood is very short (25). The short transit time is seemingly due to arrest of yeasts within small capillaries, as can occur in the ears of mice (24) and in the lumen of leptomeningeal capillaries (24, 27). Yeasts arrested in capillaries would not be detected by our methodology. In contrast, others found that blood contains both free yeast cells as well as yeast cells in the buffy coat, i.e., engulfed by phagocytic immune cells, for up to 20 days post-inoculation. Depletion of mononuclear phagocytes decreases the number of yeasts brain detected in the brain (23, 26, 34, 35). Regardless of the short transit time, there is sufficient experimental evidence to support that dissemination to the brain occurs via the hematogenous route, by direct crossing of the blood brain barrier by free yeast cells, and by indirect carriage within host mononuclear phagocytes, the Trojan-horse transport.

In models of intravenous (IV) inoculation, yeast dissemination to the brain occurs as early as 90 min after inoculation (36) (38). The speed of *C. neoformans* dissemination to the brain was attributed to the intravenous inoculation model that can simulate fungemia, an event that occurs very late in natural infections. Our findings confirm that IV infection disseminated more efficiently to the brain, but all infection routes lead to presence of viable yeasts within the brain as early as 3 h post infection. Others had detected yeasts in the brain after intranasal infection as early as 3 d (39). We did not experimentally confirm the same quick dissemination to the brain occurs with intratracheal infection, but we expect the intratracheal route of infection to be similar to the intranasal route for 2 reasons: i) at day 3 the fungal burden was similar for both routes; ii) due to sneezing (and possibly coughing reflexes), inoculation into the trachea would quickly spread *C. neoformans* throughout the entire upper respiratory tract. It is our view that once the lungs are infected, the entire upper respiratory tract becomes infected. Therefore, we believe that brain dissemination occurs very rapidly and possibly in a matter of hours, irrespective of the infection route used. This finding and others (34, 39) establish that cryptococcal brain dissemination occurs early (a matter of few hours) and simultaneously with lung infection.

After noting early dissemination post-intranasal inoculation we wondered if a route of dissemination through a nasal-olfactory system could be used by *C. neoformans*. Both *Burkholderia pseudomallei* and *Listeria monocytogenes* can reach the brain via damage to the olfactory mucosa and travelling upwards through the olfactory nerve tracts as well as other cranial nerves (28, 29). *Neisseria meningitidis* invades mouse brains by damaging the olfactory epithelium and travelling along the olfactory tract, through the cribriform plate, to invade the brain (40). The filamentous fungus *Mucor* invades the facial blood vessels from the sinus to reach the eye, the cranial nerves and the brain. In the case of *C. neoformans*, the possibility of brain dissemination through the nose has been pursued previously (41). In mice, yeast cells were observed along the olfactory nerve and the meninges starting at day 3 post intranasal instillation (41). Our study differs from this previous study because we focused on the earliest stages of infection and found no evidence of an upward invasion of the brain. We observed no signs of mucosal damage and no inflammation in the nose cavity, despite the presence of abundant yeasts and secreted GXM. In interpreting these results, we relied on the terminology of the damage-response framework that defines infection as the acquisition of the microbe by the host and colonization as a state where the damage resulting from the host-microbe interaction is insufficient to affect homeostasis (42). From this perspective, the lack of visible damage suggests that infection leads simply to nasal colonization and not overt disease. Given the lack of damage and inflammatory infiltrate our findings do not support the notion that invasion of nasal structures is involved in dissemination of *C. neoformans*. Nevertheless, intranasal inoculation results in nasal colonization where fungal cells may later interact with local immune defenses and potentially affect the development of the immune response.

Animal models show that some animals can harbor *C. neoformans* in nose for extended periods of time. Intranasal instillation resulted in yeast cells detected in the nasal cavities for periods up to 1 month of both mice and rats (43) and this infection is large enough to be detected by whole-animal non-invasive imaging as early as 1.5 weeks after infection (39). In immunocompetent laboratory-mice infected intranasally, yeasts can be detected up to 90 d after instillation (44), and guinea pigs carried *C. neoformans* in nose for several weeks (33). Some strains of *C. neoformans* are rhinotropic, with the onset of nasal lesions occurring very late in the disease in laboratory infected mice (45). Further, *C. neoformans* is frequently detected in nose of animals outside of the laboratory setting. Asymptomatic nasal carriage of *C. neoformans* has been reported in cats, dogs and koalas (46, 47). The percentage of positive animals can reach as high as 95%, as reported for feral cats in Italy (48). *C. gattii* is also quite frequent in animals, with British Columbia region having positive nasal swabs in 4% and 7% of the cat and dog population, respectively (49). In conclusion, there is evidence that *C. neoformans* can survive and even colonize upper respiratory tract of some felines and rodents, including mice, a common experimental model. This frequent colonization is associated with disease since cryptococosis is infrequent in species such as cats (50) but relatively prevalent in koalas (47). Frequent detection of *C. neoformans* in wild animals and the prolonged detection of *C. neoformans* in noses of laboratory animals shows that *C. neoformans* (and perhaps other *Cryptococcus* species) colonizes the upper airways of animals.

Serological studies indicate that humans are exposed to *C. neoformans* at an early age (7), and that *C. neoformans* can reside in the lungs in a latent form (9). Non-pathogenic species of *Cryptococcus* spp. were identified in several body sites: healthy scalps (51), in mouths of 20% of the healthy population (52), in skin of children (53), and in breast milk (54). Lung transplant patients or bronchiectasis patients frequently identify the presence of *Cryptococcus*, but no pathogenic species (55, 56). Indeed, pathogenic species of *Cryptococcus* spp. are rarely found in microbiome studies. In nose and upper respiratory tract the majority of studies (57–62) reported no *Cryptococcus* spp isolates from nasal cultures, while one study found rare *Cryptococcus* spp. (and no *C. neoformans)* by culturing nasal cavity lavages (63). There is one notable exception: high-throughput sequencing identified *Cryptococcus neoformans* as the most abundant fungal species in the middle meatus in 60% of healthy patients and 90% of chronic rhinosinusitis patients in St. Louis, Missouri, USA (64). The explanation for this striking exception is unknown and potentially may be a technical problem. Overall, *C. neoformans* is rarely recovered from healthy humans. In contrast, *C. neoformans* is frequently isolated from upper respiratory tract of some felines. This discrepancy is likely due to a combination of increased exposure of animals to environmental reservoirs of *C. neoformans* as well as immunological differences in host-species. However this is a surprising finding since animal disease has so far recapitulated human disease (13, 15, 65).

In summary, our study shows the presence of the colonization of the upper respiratory tract of *C. neoformans* in mouse models with rapid dissemination to the brain, independently of the infection route. The rapid appearance of yeast cells in different body compartments where they initiate local immune responses suggests caution in associating a particular systemic response with a specific tissue. At the very least, our findings suggest the need to revisit long held views on cryptococcal pathogenesis in animal models of infection using the most modern cellular and immunological tools.

## Acknowledgements

The authors wish to thank Oncology Tissue Services of Johns Hopkins University for the histology and use of Tissue Scanner facilities. The authors thank the Johns Hopkins Phenotyping Core and Professor Cory Brayton for the histopathological analysis.

